# An ensemble learning approach to auto-annotation for whole-brain C. elegans imaging

**DOI:** 10.1101/180430

**Authors:** S. Wu, Y. Toyoshima, M.S. Jang, M. Kanamori, T. Teramoto, Y. Iwasaki, T. Ishihara, Y. Iino, R. Yoshida

**Affiliations:** The Institute of Statistical Mathematics, 10-3 Midori-cho, Tachikawa, Tokyo, Japan; JST-CREST, Japan; The University of Tokyo, 2-11-16 Yayoi, Bunkyo-ku, Tokyo, Japan; Kyushu University, 744 Motooka, Nishi-ku, Fukuoka-shi, Fukuoka, Japan; Ibaraki University, 4-12-1 Nakanarusawa-cho, Hitachi, Ibaraki, Japan

**Keywords:** C. elegans, Whole-brain imaging, Automated annotation, Ensemble learning

## Abstract

Shifting from individual neuron analysis to whole-brain neural network analysis opens up new research opportunities for *Caenorhabditis elegans* (*C. elegans*). An automated data processing pipeline, including neuron detection, segmentation, tracking and annotation, will significantly improve the efficiency of analyzing whole-brain *C. elegans* imaging. The resulting large data sets may motivate new scientific discovery by exploiting many promising analysis tools for *big data*. In this study, we focus on the development of an automated annotation procedure. With only around 180 neurons in the central nervous system of a *C. elegans*, the annotation of each individual neuron still remains a major challenge because of the high density in space, similarity in neuron shape, unpredictable distortion of the worm’s head during motion, intrinsic variations during worm development, etc. We use an ensemble learning approach to achieve around 25% error for a test based on real experimental data. Also, we demonstrate the importance of exploring extra source of information for annotation other than the neuron positions.

## 1 Background

*Caenorhabditis elegans* (*C. elegans*) is the first living organism to have a complete genome sequencing and neural circuit mapping with all neurons named [28] thanks to its transparent body and the relatively simple neural system. Since the introduction of *C. elegans* as a model organism for neurobiology by Dr. Sydney Brenner [2], many important scientific discoveries have been achieved in the field of neuroscience, as well as biomedical research [7]. However, previous studies have focused on individual neurons only [24, 4, 11], which is not sufficient for understanding the mechanism of neural activities at the level of dynamic network. Recent advancement in bioimaging technology allows researchers to efficiently obtain high quality 4D whole-brain images of *C. elegans* [19]. For example, our laboratory can produce 15 sets of 4D images per week. Nevertheless, the average number of analyzed samples reported in the literature is only around 4 to 5 sets of 4D images [8, 18, 27]. The main bottleneck comes from the intensive demand on human effort to complete tracking and annotation for a single set of images. Therefore, there has been many studies on the automation of these data processing steps.

To achieve the ultimate goal of fully automated data processing pipeline for whole-brain images of *C. elegans*, it is important to have reliable algorithms for detecting neurons from a series of 3D images, segmenting the voxels for neuron size estimation, tracking the identified neurons along time and annotating neurons for functional analyses. Researchers have reached certain success for detection, segmentation and tracking [25, 26]. However, for annotation, existing literatures mainly focused on the body cells [12, 20, 1, 6]. The neurons of *C. elegans* are much more similar to each other and densely located in the head region that makes it extremely difficult to distinguish each of the neurons individually. This is the reason why techniques from previous success of body cell annotation or brain segmentation/registration [5] are not transferable to our case.

Annotation of neurons refers to the process of matching neurons in a new image to neurons in a reference image that has already been labeled with names, i.e., naming the unlabeled neurons in the new image. Actual images often exhibit high heterogeneity due to imaging error or changes of the worm’s posture. Therefore, the annotation task cannot be done by merely overlaying the new image on the reference image, and most of the image registration algorithms that assume small perturbations between the images would fail. To overcome such a difficulty, human annotator accumulates experience from looking through many images and searching for patterns across multiple sources of information other than the neuron position, such as neuron size and average voxel intensity. A successful automatic annotation algorithm shall possess similar abilities to handle various uncertainties in a new image by integrating different information available. In this paper, we implement such an idea through an ensemble learning approach. The details of our method is discussed in Section 2, followed by some test results and discussions.

## 2 Methods

When matching images of neurons from two distinct worms, we expect multiple sources of uncertainties, such as experimental distortion, body posture difference, and intrinsic variations during worm development. To handle these uncertainties for a more robust annotation algorithm, we propose to use the idea of ensemble learning. Our approach includes three parts:

1. Atlas generation — A large set of annotated references (atlases) are prepared to capture the variations observed in the *C. elegans*. The whole set resembles the accumulated experience/knowledge of a human annotator from many existing images.
2. Bipartite graph matching — All possible pairs of matching between neurons in a target image and a atlas is considered. Each pairs of matching is assigned a weighting value based on selected features, such as the relative distance between neurons. We optimize the total weights with constraint that enforce an 1-to-1 matching between the two images. The solution is expected to provide us a reasonable matching result for labeling the neurons in the target image.
3. Majority voting — The results of a target matched with all the atlases are combined through a simple voting system to reach a consent for the final solution. We also include a post-processing step to iteratively correct some potential mistakes by exploiting the ensemble learning method.

Figure 1 shows a summary of our auto-annotation algorithm. Details of each part are described in the following sections.

**Figure 1:**
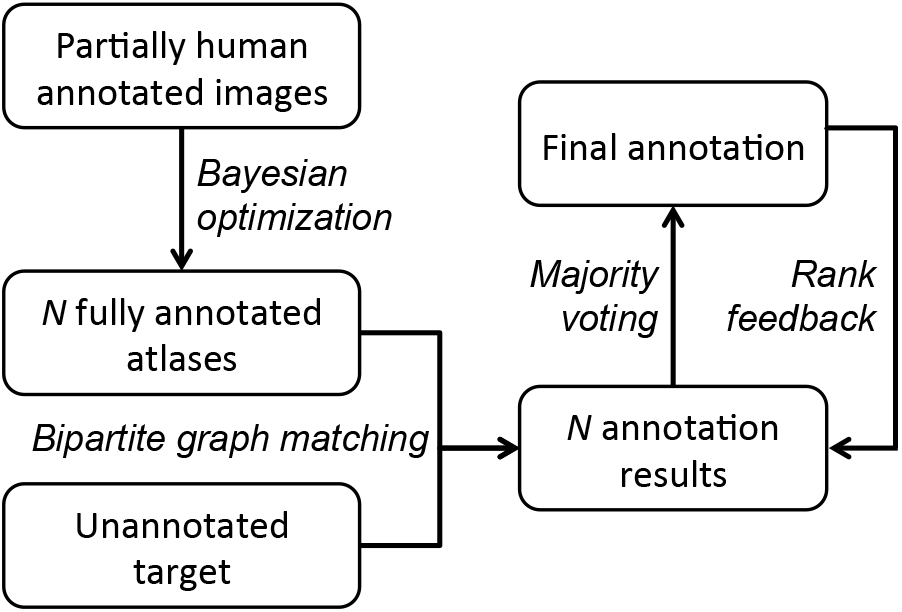
Flowchart of the auto-annotation algorithm. A summary of the algorithm that annotates a target image using partially human annotated images.

### 2.1 Atlas generation using partially human annotated data

An ideal set of references for auto-annotation would be the real 3D whole-brain images of *C. elegans* with human annotation, which come from independent worm samples. In practice, due to the difficulty and the time consuming nature of the annotation problem, we often have partially annotated data samples instead. In our study, we have a total of 190 independent worm samples annotated by human based on different promoters (a region of DNA that drives transcription of a gene, used for visualizing a subset of neurons). Appendix A illustrates the experiment details to obtain the data. The total number of detected neurons are around 190 to 210, and around 10–70% of them are annotated (see Figure 2). We combine the partial information embedded in these samples with a constraint to preserve the spatial properties of each neuron learned from the samples. The partially annotated data are randomly lined up in a series and sequentially combined through an optimization scheme to produce a single atlas. This process is controlled by a customized metric for the spatial properties of neurons and Bayesian optimization. A large set of atlases is generated by repeating the process many times.

**Figure 2:**
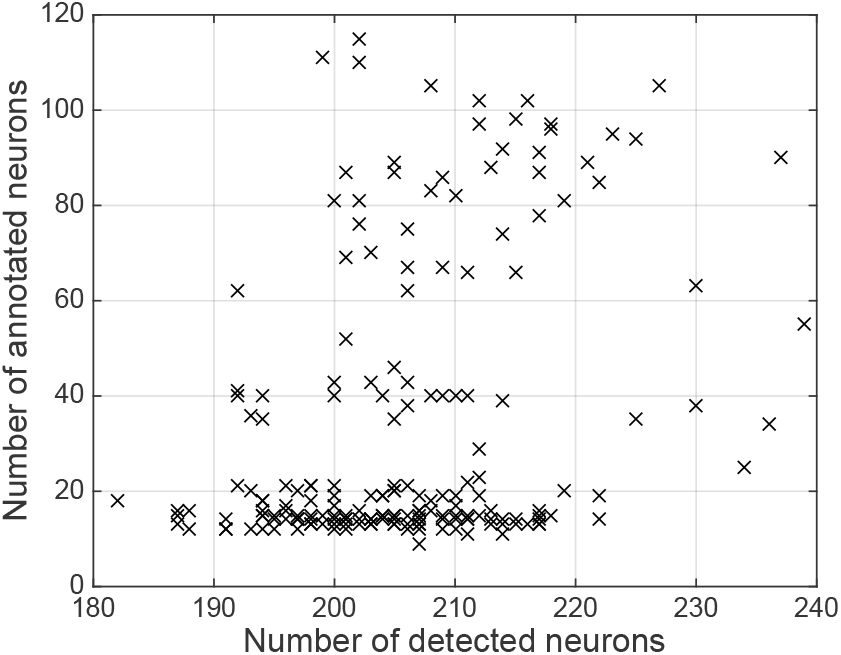
Distributions of detected and annotated neurons in the 190 images. Each cross represents one out of the 190 partially annotated samples. Around 2/3 of the samples have less than 15% of the neurons annotated.

#### 2.1.1 Customized metric

We define a metric to measure how coherent the motion of a neuron is when moving with the neighboring neurons. Consider two 3D point cloud images *I_a_* and *I_b_* of neurons (e.g., two different samples similar to Figure 13 in Appendix A), there exist *N_a,b_* neurons that are observed in both *I_a_* and *I_b_*. We would like to quantify the relative position change of all neurons. We define the displacement of each of the coexisting neurons, i.e., annotated neurons with the same name between two samples, from *I_a_* to *I_b_* to be 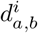, for all *i* = 1*, …, N_a,b_*. Then, we can calculate the mean displacement field 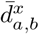 for each location ***x*** in the field of interest as follow:

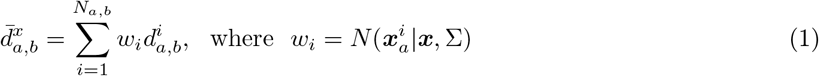

Here, *N* (*z|μ,* Σ) denotes the normal distribution value at *z* with mean *μ* and covariance matrix Σ, and 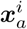 denotes the location of the *i^th^* coexisting neuron in *I_a_*. Conceptually, 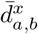 represents a spatial smoothing of the *N_a,b_* displacement samples based on a decaying weight function, i.e., 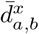 is the average movement of other neurons in the perspective of neuron at *x* (movement of the closer neurons is more important than the further ones). We consider a *coherent motion* to be an observed displacement of a neuron that follows the mean displacement field 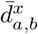, i.e., to follow the trend around yourself.

#### 2.1.2 Generate atlases by Bayesian optimization

Based on our notion of coherent motion, we apply the Bayesian optimization algorithm to generate atlases such that the coherency level of each neuron in the set of atlases is statistically similar to the one in the human annotated data set. Bayesian optimization is a stochastic optimization method aiming at global optimization [13]. We will begin with the description of the atlas generation algorithm with parameters to control the level of coherency imposed during the combination of partially annotated data. Then, we illustrate the use of Bayesian optimization to determine the parameter values by minimizing the difference of the coherency statistics between the generated atlas set and the actual data set.

To generate one atlas, we take the following steps (see Figure 3):

1. Generate a randomly ordered sequence of the partially annotated data set.
2. Sequentially combine an image data *I_b_* to the next one *I_a_*:

a. All annotated neurons in *I_b_* remains the same.
b. All annotated neurons that coexist in both *I_b_* and *I_a_* are used as samples to construct the mean displacement field 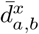 for a pre-determined Σ value in equation 1.
c. All annotated neurons in *I_a_* but not in *I_b_* are moved according to 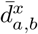.

**Figure 3:**
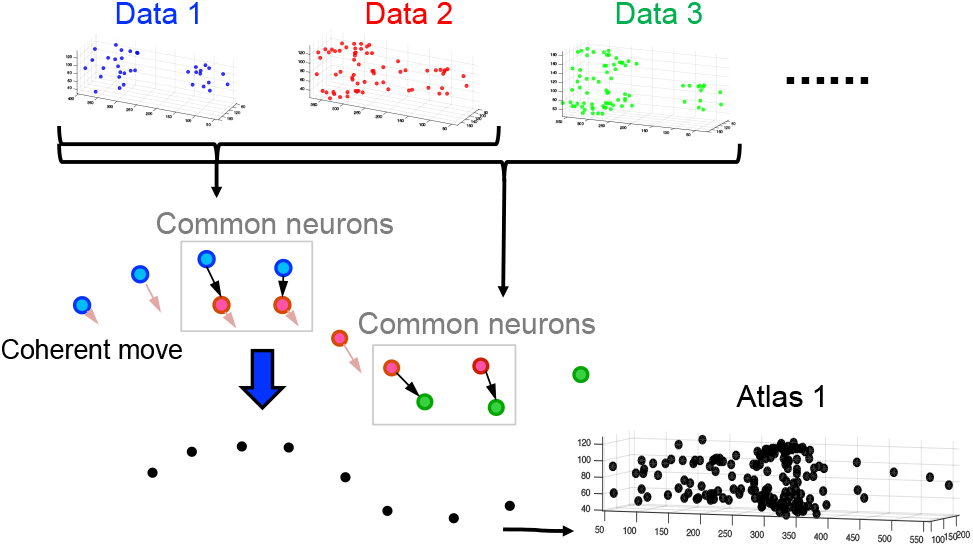
Combining partially annotated data to one atlas. We reorder the data set to a random sequence and merge them one by one based on the notion of coherent motion.

In this process, configuration of the neurons will be dominated by the early chosen data in the random sequence. To alleviate the effect of noise, we skip the combination of *I_a_* and *I_b_* that shares less than 5 neurons in common. At the end, we generate a large set of atlases by creating many random sequences under a chosen parameter value for Σ, which is the covariance matrix defining the strength of the spatial smoothing for the mean displacement fields.

Next, we define a metric for motion coherency 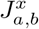 for the *i^th^* neuron in an image *I_a_* that coexists in an image *I_b_* as:

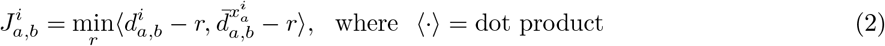

This metric measures the difference between the observed displacement of a neuron and its mean displacement field up to a constant shift. We calculate 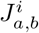 for all neurons in all possible pairs of *I_a_* and *I_b_* in the partially annotated data set. We can then evaluate the mean and standard deviation of the metric value for each neuron observed in the data set, which defines a Gaussian distribution. We perform the same calculation for the resulting atlas set, and use Bayesian optimization to minimize the Kullback-Liebler divergence between the Gaussian distributions for all neurons in the partially annotated data set and in the atlas set. During the optimization, we ignore the neurons that coexist in less than a total of 5 image data to avoid sensitivity to noise. We have used the R-package of Bayesian optimization implementation in our study [29].

### 2.2 Bipartite graph matching with a pre-processing step

The core of our annotation algorithm is to match a neuron in a target unannotated image to a neuron in an annotated reference image (an atlas). To account for the body posture difference between the worm in the target image and the atlas, we pre-processed the reference image by applying a non-rigid registration algorithm, called the Coherent Point Drift (CPD) [16, 15]. Then, we framed the neuron annotation problem as a bipartite graph matching problem, solving it with the well-known Hungarian algorithm [10, 14].

#### 2.2.1 CPD pre-processing

From our experience, even when the *C. elegans* is well constrained in a small tube during experiment, its head can still undergo noticeable movements. The resulting images from different worms may be coming from very different body posture, as well as the variation due to the different worm size. These differences between the images can be seen as undergoing some highly non-linear deformations. A robust non-rigid registration is needed to guarantee a good annotation result. An important assumption we can made in this case is the notion of motion coherency. This is equivalent to the assumption that the relative position between neurons shall be roughly maintained across different images. The Motion Coherence Theory (MCT) provides an explicit mathematical formulation for this purpose [30, 31]. We use the Coherent Point Drift (CPD) algorithm, which is an efficient image registration implementation of the MCT idea [16, 15], as a pre-processing step to increase the matching accuracy. We first apply the CPD algorithm to an atlas, and then perform the matching of neurons between the two images to achieve auto-annotation. We note that [16, 15] have shown that CPD generally works better than other affine transformation methods, or even other non-rigid transformation algorithms, when the mapping between the two images are highly non-linear. Figure 4 shows an example of the effect of CPD on our data sets. In our study, we used the MATLAB implementation of the CPD provided in [16, 15], and fixed the two important parameters *β* = 2 and λ = 3 based on the suggestions in the paper and our empirical tests.

**Figure 4:**
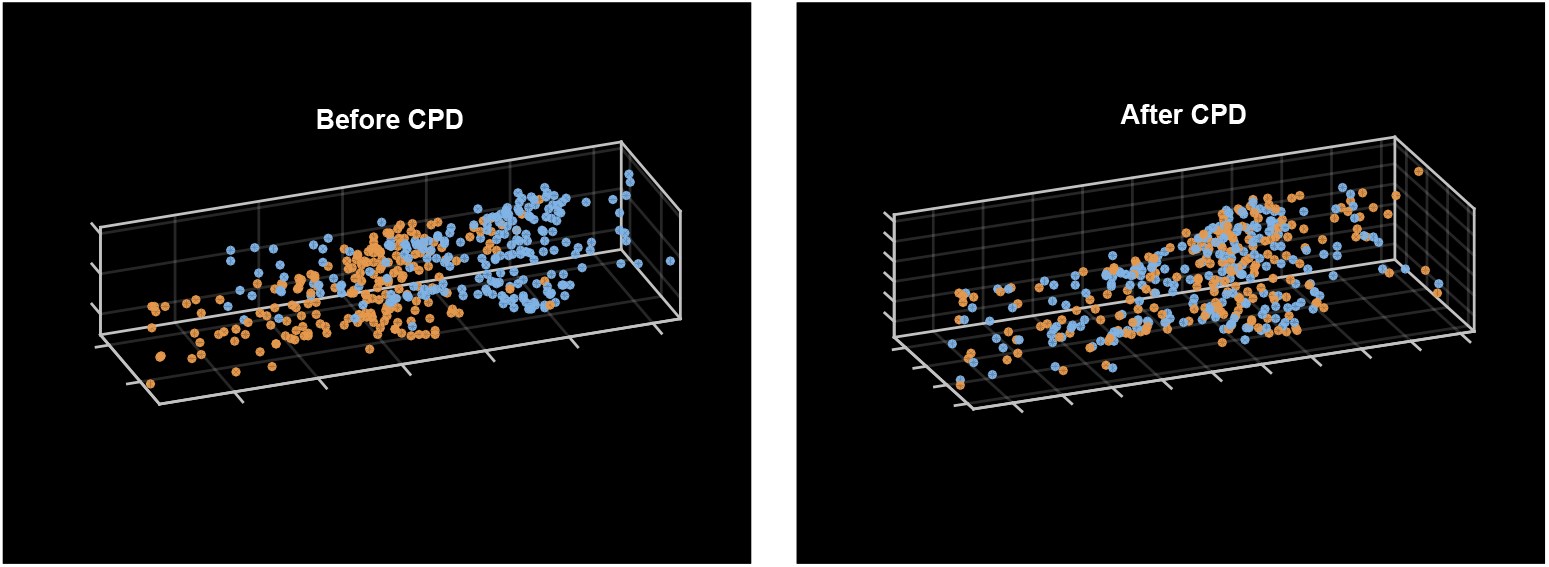
Example of using CPD to register two images. Orange dots represent the target data set, which remains unchanged. Blue dots represent the atlas, which CPD is applied on to match with the target data set.

#### 2.2.2 Bipartite graph matching

After the pre-processing step, neurons in the target image and an atlas are matched, which can be seen as a bipartite graph matching problem. The matching is achieved by comparing one or more selected features between neurons. Here, features refer to some quantitative properties for the neurons that can be used to distinguish the identity of a neuron from another. For example, the most fundamental feature for point cloud registration problems is the spatial distance between two points in a reference space. In our case, this is simply the spatial distance between any two neurons in the target data and the atlas after CPD pre-processing. Other features typically considered for cell or neuron matching include cell/neuron volume, brightness of the signal captured in the image, etc. For our auto-annotation algorithm, each feature is represented by a two dimensional matrix *A*, where the {*i, j*} entry in the matrix is the feature value calculated from the *i*^th^ neuron in the target data to the *j*^th^ neuron in the atlas. When there are *N_f_* features chosen, we assemble them into a single matrix *A*_BGM_:

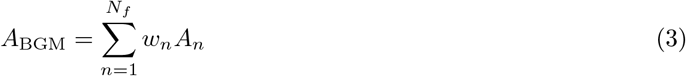

where *A_n_* is the matrix representing the *n*^th^ feature. For a given set of matchings between the neurons in the two data sets, where each neuron is only allowed to be matched with one neuron, we can calculate the sum of the values in *A*_BGM_ that correspond to the selected matchings. This value represents the likelihood of the given matching set. By maximizing or minimizing (depending on the meaning of the feature values) the likelihood value, we can obtain the final matching of the neurons between the target data and the atlas. Such an optimization problem can be done by the well-known Hungarian algorithm [10, 14].

In our test shown in Section 3, we have chosen only the basic features as preliminary test: the spatial distance between neurons. We have tried to use other features mentioned above as well. At the end, we did not choose the features related to neuron size or brightness in the image because our experimental results indicated that these information is too noisy to be used in our case. In the case of more than one selected features, the weight *w_n_* for each feature is determined by the classical statistical method, data splitting, i.e., we can separate our data into two sets and use one of them to optimize for *w_n_*.

### 2.3 Majority voting with “rank-feedback"

To handle the uncertain variations of the target images in practice, we use the majority voting approach for a more robust matching solution. Assuming our atlases are capturing the variations well, we match a target image without annotation using *N_a_* atlases to produce *N_a_* sets of matching results. Each result is considered as one vote towards a possible matching pair. We collect all the votes to obtain a final vote table that represents the level of confidence for each matching pair. To obtain the final solution, we can simply apply the Hungarian algorithm to this vote table to optimize for the votes. The overall procedure is outlined in Figure 5. On the other hand, the ranking of different possible matching for a neuron and the vote margins contain valuable information about the uncertainty of the matching. We can exploit the ranking information to further improve the matching/annotation results using the idea of “rank-feedback", described in the following.

**Figure 5:**
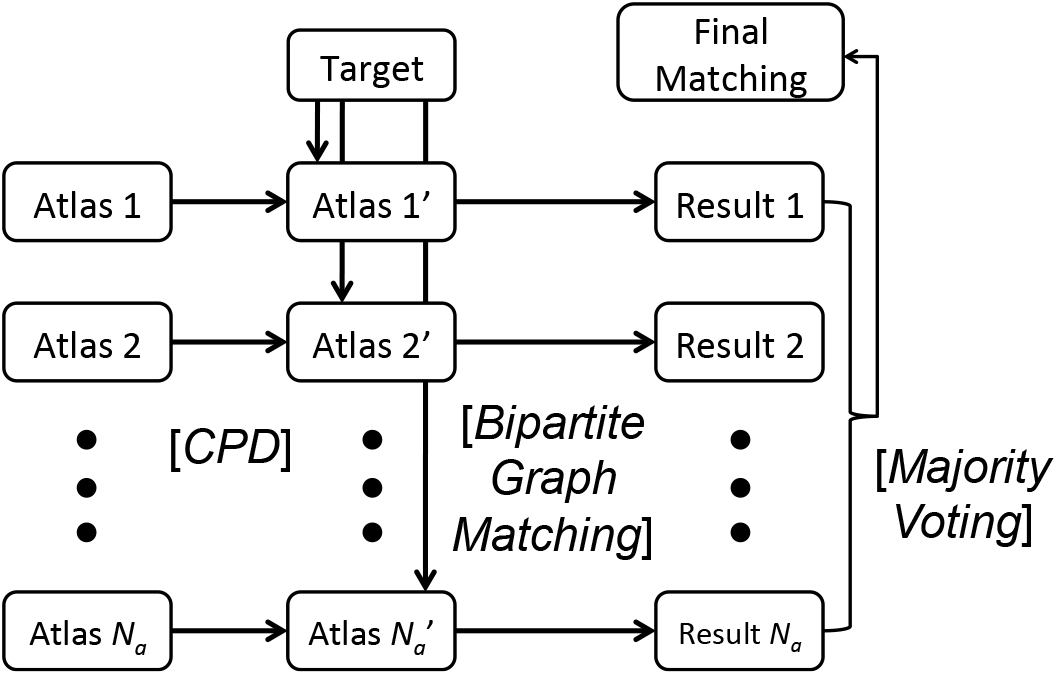
Outline of the matching procedure for auto-annotation. A summary of the algorithm that matches neurons from a target image to different neuron names. Here, neurons in the target image are unannotated, and neurons in all atlases are annotated.

After performing one round of majority voting, the top *N_r_* matchings represent a cluster of neurons that is similar to a targeted unknown neuron. This information can be re-used as a new feature added to the bipartite graph matching. For example, the links corresponding to the top *N_r_* matchings will have feature value 1, and 0 otherwise. By re-doing the same matching procedure with this extra feature, this allow us to focus on matching the more probable pairs in the next round of majority voting. As we decrease the number *N_r_*, we are focusing on a smaller and smaller cluster of possible solutions. Conceptually, this is similar to the *coarse-to-fine* approach in the typical image registration problem (see Figure 6). We call this idea the “rank-feedback" method for majority voting, and it can be summarized as the following steps:

1. Perform matching with majority voting to the target using pre-defined atlases.
2. Record the top *X* voting results (Rank *X*) for each target neuron required to be named.
3. Repeat step 1 with the top *X* voting information included in the bipartite graph matching algorithm as an extra feature, stop after convergence.

For actual implementation, we suggest the following based on our experience:

1. The selected value for *X* should be decreased for each iteration of the convergence loop.
2. The loop shall be stopped with *X* = 3.
3. Information of the top *X* voting from the previous steps in the loop shall be used accumulatively, i.e., all of the feature values obtained during the loop shall be added up.

**Figure 6:**
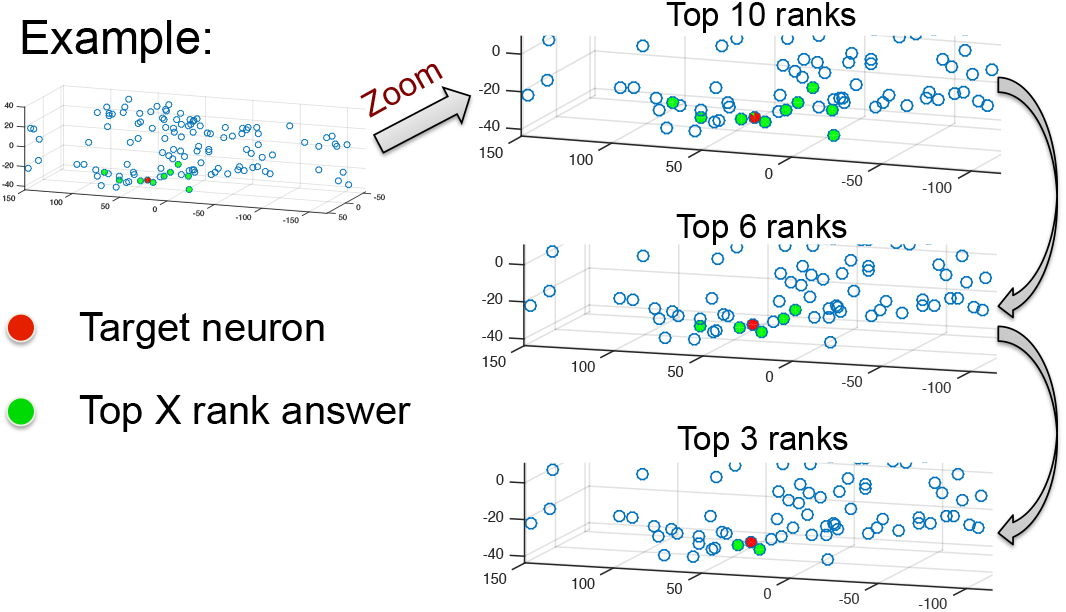
Coarse-to-fine effect of rank-feedback majority voting. When grouping the top *X* possible matches for each neuron with decreasing *X*, it allows the possible matching solution to gradually converge. This approach is similar to the tempering idea often used in optimization problems. Overall, this is similar to a typical coarse-to-fine approach in image registration problems.

## 3 Results

We performed two tests to evaluate the performance of our method. The first test is simply using the atlases we generated to be the target test set. In this case, the assumption that the atlases capture well the uncertain variations of the target data is trivially satisfied. Hence, a good result is expected. The second test involves test data coming from a different laboratory, thus, a more realistic and challenging test.

### 3.1 Simple test with generated atlases as target

#### 3.1.1 Data preparation

In the first test, we randomly picked 200 atlases out of around 60000 atlases that we have generated based on the procedure mentioned in the Section 2. These atlases are treated as unannotated targets, and the neuron names recorded in our database provide the final solution for testing the algorithm accuracy. We randomly selected 1000 atlases from our full atlas set as references that are different from the test data in order to perform the majority voting procedure in our method.

#### 3.1.2 Test results

First, we perform bipartite graph matching to match each of the target data in the test set with the 1000 atlases according to the details explained in the method section. We obtain a total of 200000 sets of matching results. Then, for each of the 200 targets, we combine the 1000 matching results using the majority voting approach. The results are summarized in Figure 7. Using only the basic bipartite graph matching method results in ~40% error rate on average (refer to the histogram in the figure). Simply adding the majority voting method without rank-feedback already leads us to less than 10% error rate. If we consider the accuracy for the top 3 voted matchings, we achieve less than 1% error rate. The rank-feedback has limited margin for improvements, thus, we directly moved on to the realistic test.

**Figure 7:**
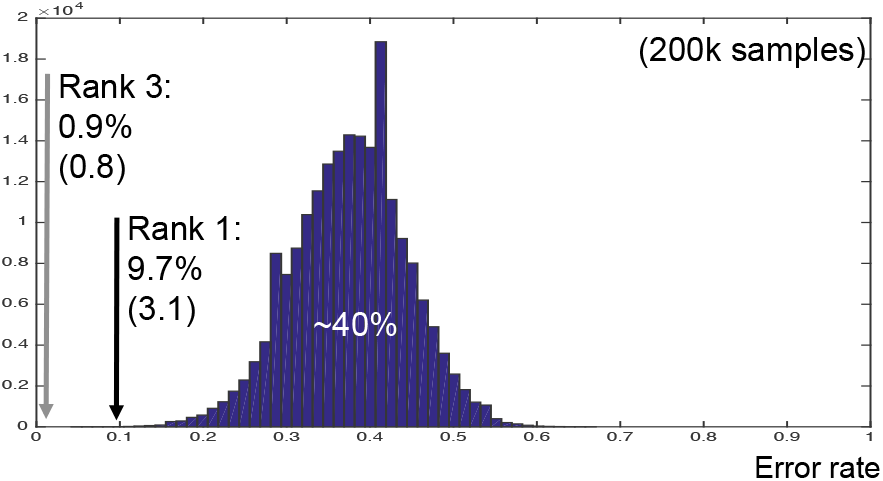
Error rates of auto-annotation for the first test. The histogram shows results of annotation without majority voting (200000 samples), with a mean error rate at around 40%. The solid arrows indicate results of majority voting without rank-feedback. Standard deviations of the majority voting results from the 200 samples are shown in parentheses. *Rank X* indicates results based on the top *X* voting, i.e., the matching is considered correct if any matching within the top *X* voting is correct.

### 3.2 Realistic test with new target data

#### 3.2.1 Data preparation

In the second test, we aim for a more realistic validation for our algorithm. We created a separate data set from a different laboratory with different experimental settings, but the basic preparations of the worm samples remain the same as those used for atlas generation. A worm was placed under a customized spinning disk confocal microscopy and fluorescence images were obtained in three channels, CFP, YFP, and mCherry, simultaneously. The sizes of the image along the *x* and *y* axes were 512 and 256, respectively, and the sizes of a voxel along the *x* and *y* were 0.33 *μ*m. Along the *z* axis, the image stacks covered 28 *μ*m range (a total of 20 stacks). We use a total of 94 frames as test target which were annotated and inspected by human. There are a total of 205 detected neurons, in which only 134 of them are manually annotated. Hence, we performed the test based on those neurons only. Figure 8 shows an example of the raw images used in this test with *roiedit3D* processing.

**Figure 8:**
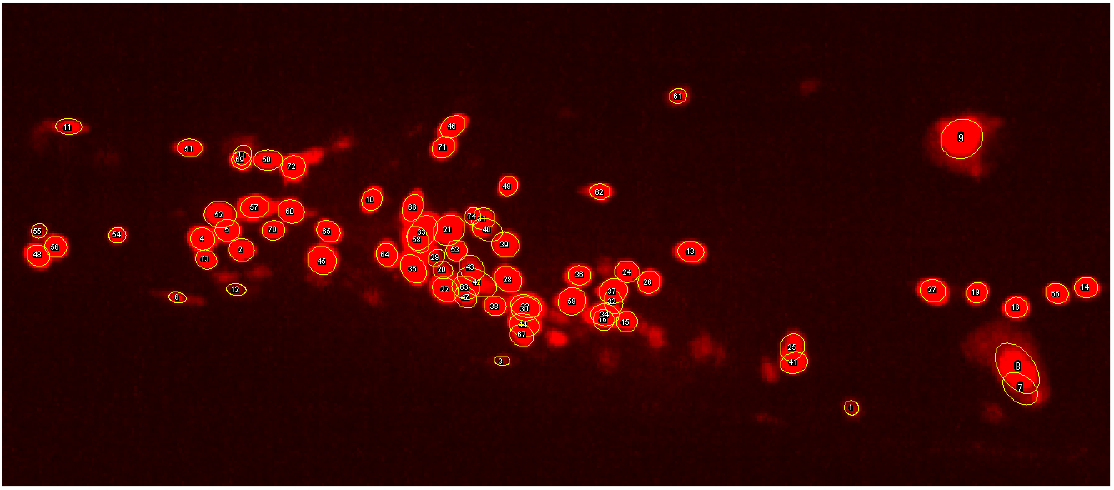
Example of unannotated target image after detection and tracking. Pan-neural images of C. elegans with mCherry promoter (in red). Yellow circles are the detected neurons using *roiedit3D* with temporary numbers labeled.

#### 3.2.2 Test results

First, we perform bipartite graph matching to match each of the target images in the test set with the 1000 atlases according to the details explained in the Section 2. We obtain a total of 94000 sets of matching results. Then, for each of the 94 targets, we combine the 1000 matching results using the majority voting approach. The results are summarized in Figure 9. By using the basic majority voting, we achieve an immediate ~20% improvement, which bring us almost to the best possible matching cases within the 94000 matching cases. However, the error rate is still around 40%, which is similar to the worst case of the first test. We note that there is a significant improvement of ~25% if we consider the top 3 voted matchings. Hence, we applied the rank-feedback idea to exploit this situation, and indeed, it gives us another ~10% improvement.

**Figure 9:**
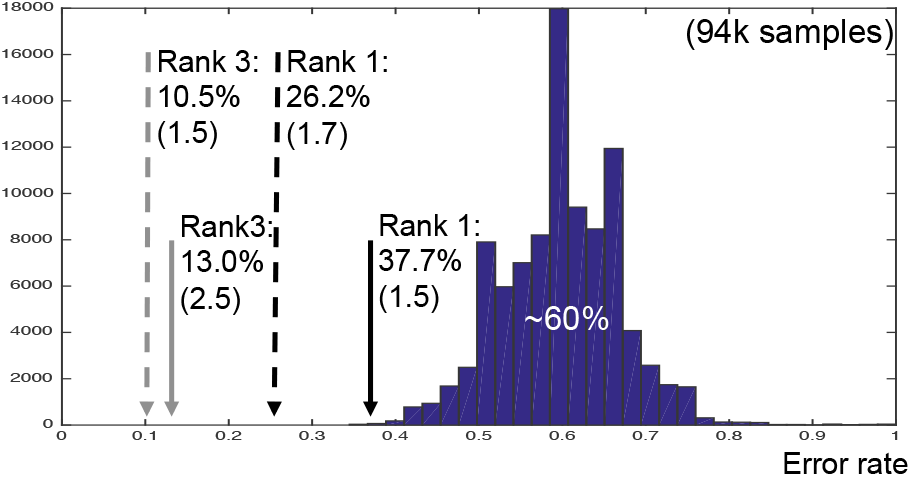
Error rates of auto-annotation for the second test. The histogram shows results of annotation without majority voting (94000 samples), with a mean error rate at around 60%. The solid arrows indicate results of majority voting without rank-feedback. The dashed arrows indicate results of majority voting with rank-feedback. Standard deviations of the majority voting results from the 94 samples are shown in parentheses. *Rank X* indicates results based on the top *X* voting, i.e., the matching is considered correct if any matching within the top *X* voting is correct.

## 4 Discussion

The test data in the first test are from the same atlas pool of the references used for auto-annotation. Hence, the references are expected to capture the features in the test set well, leading to a good annotation result. However, in the second test, the test set is expected to be very different from the references, thus, resulting in an expected performance reduction. To better understand the difficulty in this matching problem, we checked the annotation accuracy for each neuron. Figure 10 shows the error rate of each neurons plotted with the neuron position. We observe that the errors mainly occur in the center part, where neurons are densely collided to each other. In those area, only position information is expected to be not sufficient for perfectly distinguishing between the neurons. Other reliable source of information is needed, for instance, utilizing an extra stable promoter as a new feature for matching. We are currently investigating the technical feasibility of this idea.

**Figure 10:**
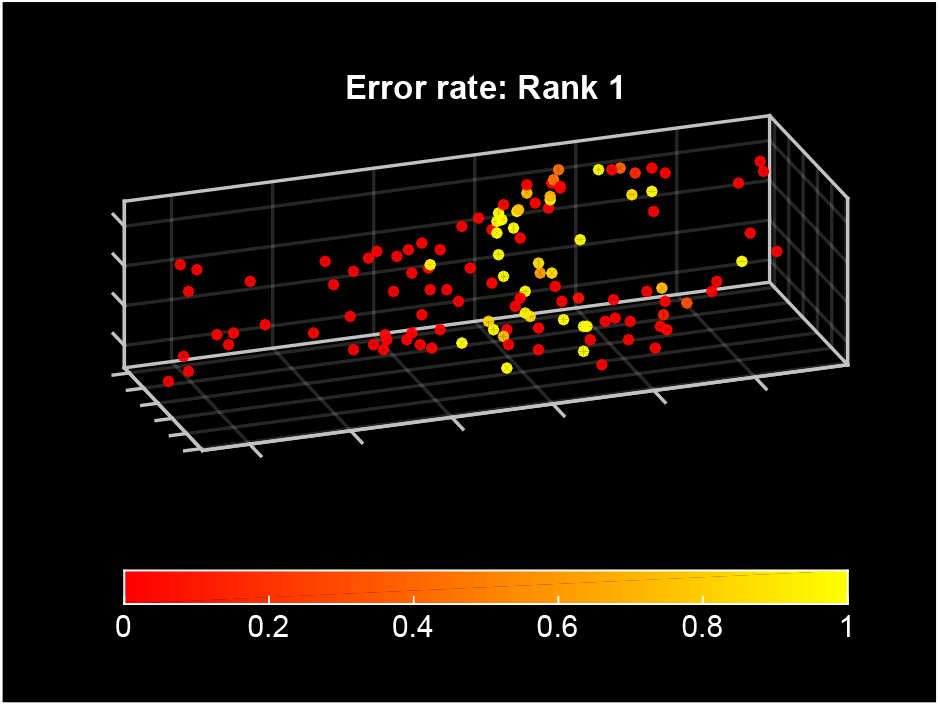
Mean error rate of each neuron. Taking one of the target data in the second test as reference for the neuron location, this figure shows the mean error rate of each neuron using a color map from red to yellow, representing zero error to always having error, respectively.

One may suspect that the bad results in the second test could be deal to insufficient number of atlases used in the majority voting step. To demonstrate that this is not true, we repeated the second test for one random target image with different number of atlases used in the majority voting stage. Rank-feedback is not used in this case. The number of atlases varies from 10 to 1000, and for each chosen number, we repeated the test for 20 times. Figure 11 shows the resulting error rate with one standard deviation error bar over the 20 trials. There is clear convergence for both the mean and the standard deviation of the error rate, thus, the number of atlases used in the majority voting step is sufficient. Furthermore, our results demonstrate the importance of using the uncertainty information obtained from the majority voting approach for further improvement (using the idea of rank-feedback). In fact, this information can also be used to assist human inspection in the future, as the top 3 voted matchings can be suggested to the human inspector as candidates of actual solutions.

**Figure 11:**
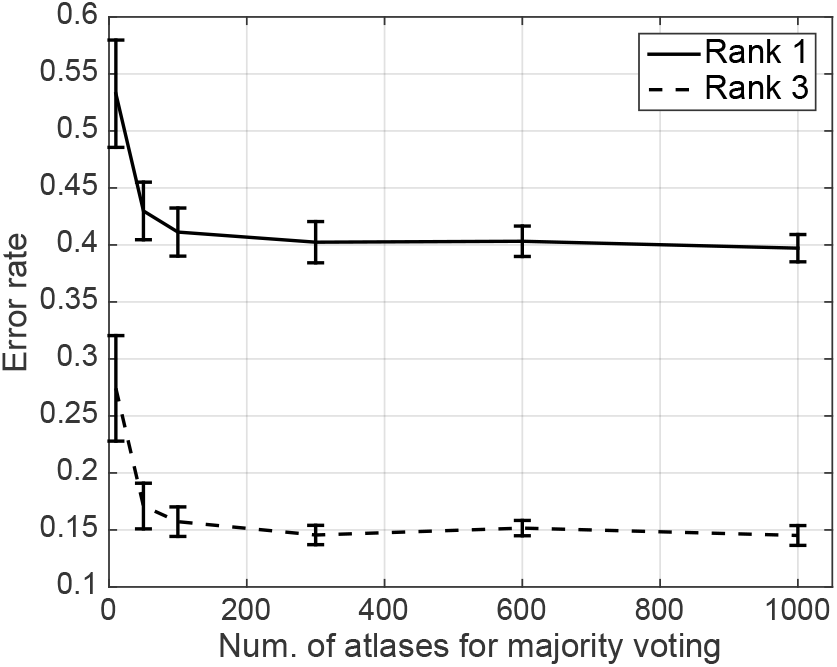
Convergence of annotation accuracy with respect to the number of atlases used for majority voting. We repeated the second test for one random target image with different number of atlases used in the majority voting stage. Rank-feedback is not used. The number of atlases varies from 10 to 1000, and for each chosen number, we repeated the test 20 times. The solid line shows the mean error rate for Rank 1 with an error bar of one standard deviation. The dashed line shows the mean error rate for Rank 3 with an error bar of one standard.

**Figure 12:**
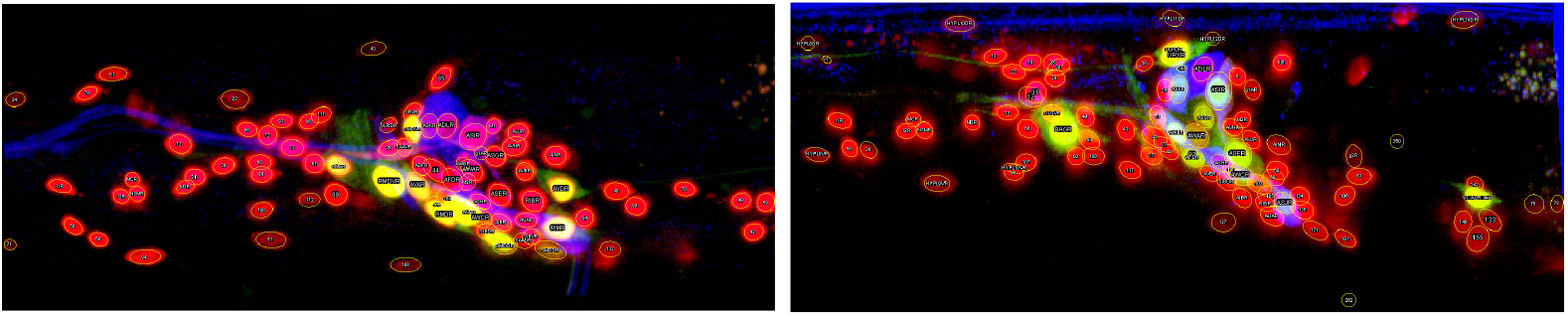
Two examples of raw image with partial human annotation using two types of promoters. Red: mCherry, green: YC2.60 expression driven by glr-1 promoter (upper) or tax-4 promoter (lower), blue: lipophilic dye, yellow circles: detected neurons using *roiedit3D* with labels (temporary numbers or annotated neuron names).

**Figure 13:**
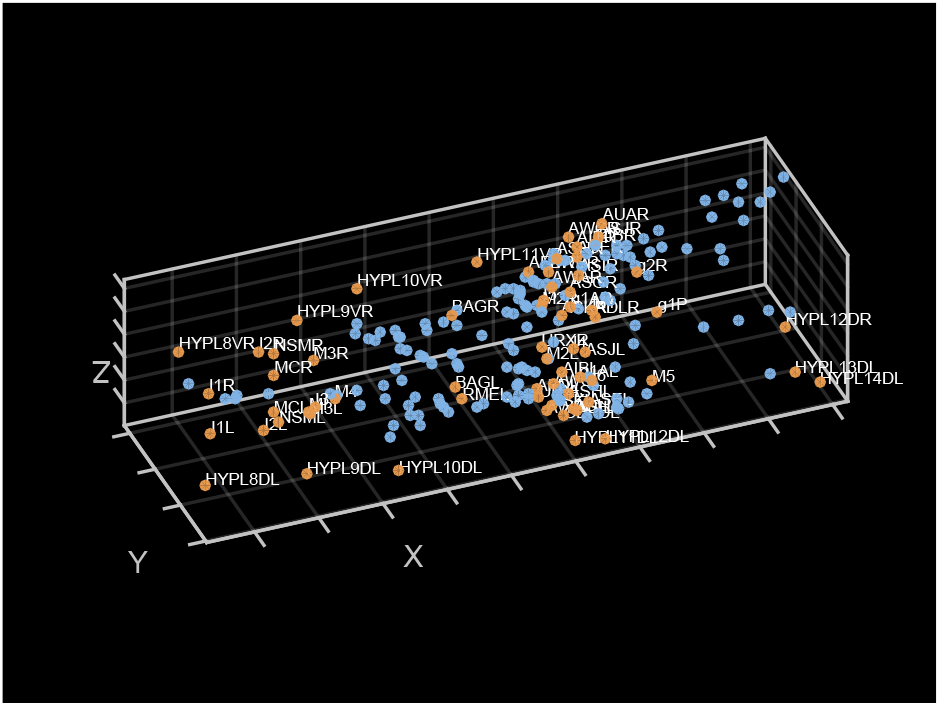
An example of partially annotated sample. Orange dots are annotated neurons with names in white, and blue dots are unannotated neurons. There are 66 annotated neurons out of a total of 211 detected neurons in this sample.

## 5 Conclusion

Our proposed auto-annotation approach based on ensemble learning method achieves around 75% accuracy in our realistic test without human correction. We expect further improvement when extra promoters are used to bring more information, which is especially important to identify neurons in dense region. We are considering this idea in our next study. We note that our current approach relies on the assumption that the atlases are good representation of the unknown target images. This assumption fails when the number of coexisting neurons within the atlases and the target image is not high. Our experience indicates that the method will fail when the difference between the number of neurons in the atlas and the target image is more than 10%. We are currently looking for robust criteria to handle such a *missing data* problem. As of now, human inspection seems to be inevitable. In our on-going work, we aim to involve the uncertainty information obtained from our auto-annotation algorithm to assist human inspection. We believe that such a semi-automatic approach will be a good solution to achieve high efficiency data analysis pipline for whole-brain images of *C. elegans* in the very near future.

## Appendix

### A Partially annotated data

To obtain the partially annotated data set, we created 3D whole-brain images based on the procedure shown in [26]. In *C. elegans*, various cell-specific promoters that drive gene expression in specific subset of neurons were identified. We generated many strains where the specific neurons were marked up with fluorescent proteins by using the cell-specific promoters [26]. The nuclei of all neurons are marked by the red fluorescent protein mCherry [23]. The nuclei of specific 22 sensory neurons are marked by Yellow-Cameleon 2.60 (YC2.60) [17] with *tax-4* promoter [9]. Other promoters used include *eat-4* [21] and *glr-1* [3]. Lipophilic dye DiO (D275, Thermo Fisher Scientific) or DiR (D12731, Thermo Fisher Scientific) [22], which stains specific 12 sensory neurons, is also used. We took volumetric still images of adult worms by laser scanning confocal microscopy (Leica SP5 with 63× water immersion lens and 2× zoom). Typical sizes of the images along the *x*, *y*, and *z* axes were 512, 256 and 170 voxels, respectively. The sizes of a voxel along these axes were 0.240, 0.240, and 0.252 *μ*m, respectively. The neurons in the images were detected, and the marked neurons were identified using *roiedit3D* [26] with human inspection. *roiedit3D* is an open-source MATLAB package that includes a very effective automatic neuron detection, segmentation and tracking algorithm with a user interface for human inspection and correction of the results. The positions, sizes and intensities of the neurons were then identified and used for human annotation. Figure 12 shows two examples of the raw images with *roiedit3D* processing and figure 13 shows an example of the post-processed partially annotated data set in the form of point clouds, which is the form of data used for atlas generation.

## Notes

* This work was supported by the JST-CREST.

